# Nanopore duplex sequencing reveals patterns of asymmetric states of 5hmC and 5mC in the medaka brain genome

**DOI:** 10.1101/2025.06.24.661039

**Authors:** Walter Santana-Garcia, Tomas Fitzgerald, Felix Loosli, Joachim Wittbrodt, Ewan Birney

## Abstract

The nucleotide modification 5-methylcytosine (5mC), has been extensively described in terms of its genomic distribution and function. However, the distribution and functional contributions of its oxidised state, 5-hydroxymethylcytosine (5hmC), and its interplay with 5mC remain to be further characterised. Moreover, the hemi-methylation and hemi-hydroxymethylation of CpG dinucleotides in the genome also remain largely unexplored. Here, we use Nanopore duplex sequencing, *i*.*e*, the sequencing of both strands of DNA from a single double stranded molecule, to profile the co-occurrence of 5hmC with 5mC in CpG dinucleotides of medaka (japanese rice paddy fish, *Oryzias latipes*) brain tissues. We characterised asymmetrical patterns of hemi-modified states of CpGs and show using duplex resolution that in the asymmetrical state 5hmC/5mC, 5hmC is preferentially found in the antisense strand of DNA in intron regions, in particular in splice sites, whereas the exons have an enrichment in 5hmC in the sense strand. Furthermore, we found that in the 5hmC/5mC state, 5hmC preferentially occurs along one strand of the DNA duplex in specific families of transposable elements. Altogether these results shed more light on understanding the function and distribution of 5hmC and its mutual role with 5mC in the genome.

## INTRODUCTION

The four canonical nucleotides in DNA can be covalently modified (Sood et al., 2019), and they can contribute to the regulation of biological processes across the tree of life. The methylation of the fifth carbon in the cytosine ring (5mC) is the most studied DNA modification (Boyes & Bird, 1991), and in vertebrate genomes, 5mC exists mostly in symmetrical pairs at the CpG dinucleotide context (*i*.*e*. one 5mC on each strand of the DNA duplex molecule). In vertebrate genomes, around 70–80% of CpG sites are methylated (Feng et al., 2010). 5mC has been historically linked to a locus-repressive role, for instance, it is normally found in permanently repressed DNA regions, such as imprinted genes and transposable elements (Bourc’his & Bestor, 2004).

Two molecular mechanisms link DNA methylation with its genomic functions: the recruitment of chromatin complexes, and the modification of binding affinity of proteins to DNA (reviewed in Greenberg & Bourc’his, 2019). For instance, methylated cytosines in the CpG dinucleotide context can recruit histone deacetylase (HDAC)-containing complexes that promote transcriptional repression in the locus where methylated CpGs are present (Jones et al., 1998). 5mC can also tune the DNA-binding of transcription factors (TFs), as some TF families have lower DNA affinity when 5mC are deposited in their binding sites, whilst others such as some homeodomain proteins increase their DNA binding (Yin et al., 2017).

The ubiquitous functions of 5mC in the regulation of molecular processes in a context-dependent manner, suggests a dynamic and flexible nature of 5mC in the genome. In fact, it was recently demonstrated that even the methylation pattern of a CpG, traditionally regarded as symmetrical, can be broken (C. Xu & Corces, 2018) to produce persistent hemi-methylated states (*i*.*e*. the methylation of only one strand of a CpG). In addition, Xu & Corces, 2018, showed that hemi-methylation patterns can not only be stably inherited through several cell divisions, but that they have an active role in increasing chromatin interaction with CCCTC-binding factor (CTCF) and subsequently affect genomic contact patterns.

5mC can itself be modified, as it can be progressively oxidised to 5-hydroxymethylcytosine (5hmC), 5-formylcytosine (5fC) and 5-carboxylcytosine (5caC), by the iterative oxidizing catalysis of the ten eleven translocation (TET) enzymes. The DNA repair machinery can then restore the base to an unmodified cytosine. This active demethylation pathway is present in all cells and considered in dynamic equilibrium with the methylation pathways. Among the oxidized states, 5hmC is a more stable and abundant modification in the genome, particularly enriched in post-mitotic neural cells (Diotel et al., 2017). 5hmC has also been associated with gene transcription, as it is more present in gene bodies of transcribed genes and the TET1 enzyme is found at transcription start sites (TSS; Huang et al., 2014). Additionally, the dysregulation of 5hmC in the genome might have a role in cancer, as 5hmC levels are lowered in many cancer types, co-occurring with mutated TET proteins (Jin et al., 2012). Although the simplest explanation for the occurrence of 5hmC is that it is a transient oxidative state of 5mC, the functional contributions of 5hmC and its interplay with 5mC, particularly in CpG contexts are still largely unexplored. In this paper we will use the term 5mC to mean methylated cytosine, 5hmC hydroxymethylated cytosine and 5C as unmodified, canonical, cytosine.

The reliable and simultaneous detection of 5mC and 5hmC in DNA is important to characterize their mutual roles in the genome. Although useful, traditional experiments used to detect 5mC and 5hmC, such as Whole-Genome Bisulfite Sequencing (WGBS; Lister et al., 2009) and TAB-seq (Yu et al., 2012) can be difficult to implement, in particular for 5hmC, and the simultaneous profiling of 5mC and 5hmC in the two strands of CpGs at the same DNA duplex molecule is not possible. Importantly a single bisulfite reaction will convert both 5mC and 5hmC, meaning that WGBS based read out is the union of these two modifications. Recently, Oxford Nanopore Technologies (ONT) enabled the sequencing and simultaneous profiling of different DNA modifications, including 5mC and 5hmC. In addition, ONT sequencing can sequence both strands of the same DNA molecule, termed duplex sequencing. Duplex sequence occurs when a double stranded molecule is ligated to an ONT motor at both ends, and having sequenced one strand the second strand follows immediately through the same nanopore. This pairing of molecules can be recognised by the precise matching of the molecule start and end, and the timing of the second strand through the pore.

In this study, we exploited the presence of nanopore sequencing for a large panel of medaka fish which was performed to provide accurate de novo assemblies. However, the preparation method provided substantial levels of duplex molecules, meaning that we have high coverage of nanopore based duplex molecules and serendipitously we used neuronal tissues as a source of DNA. This provides to our knowledge the largest duplex molecule dataset of a vertebrate genome derived from neuronal tissue. We are able to comprehensively profile the presence of 5mC and 5hmC in a duplex context, allowing us to assess hemi-methylation and hemi-hydroxymethylation at large scale for the first time.

## RESULTS

In order to profile the co-occurrence patterns of 5mC, 5hmC and 5C in both strands of CpG dinucleotides, we sequenced the brain genome of 50 medaka individuals, using ONT sequencing (Figure 1) from the Medaka Inbred Kiyosu-Karlsruhe (MIKK) panel (Fitzgerald et al., 2022). We ensured we basecalled the reads with a 5mC and 5hmC model and with duplex detection enabled. On average we obtained 18.9% of simplex (single pass) reads in a duplex form, *i*.*e*. 9.45% of molecules sequenced were sequenced in both strands. (Supplementary Figure 1). Over the 50 samples this resulted in a 205X mean coverage of duplex form sequencing across the medaka genome.

**Figure 1.**
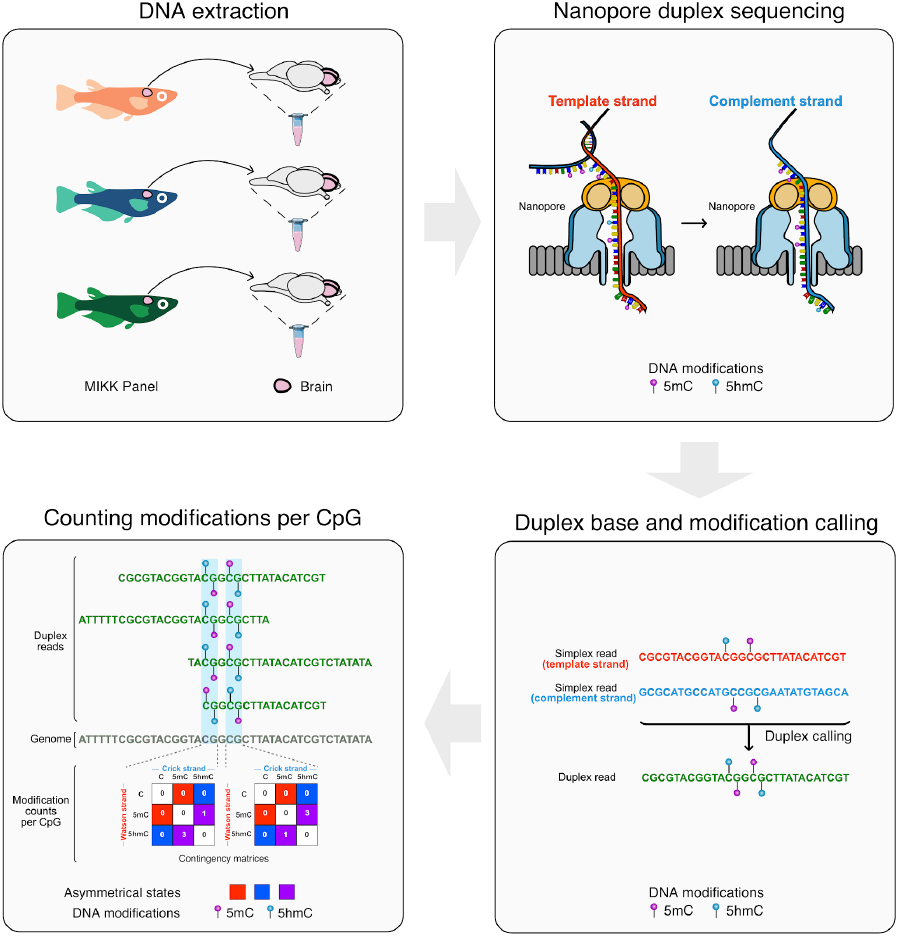
Schematic workflow of study design. The processing of medaka samples for DNA extraction, sequencing, duplex read calling and counting strategy of modification patterns on CpG is depicted.

After base and modification calling duplex reads in the 50 MIKK individuals, we assigned the CpGs in duplex reads to one of nine possible states and quantified their occurrence (Figure 1). From this quantification, we confirmed that when CpGs are methylated in the genome they are predominantly symmetrically methylated (5mC/5mC) in both strands (Figure 2). The extent of this modification pattern when one considers that WGBS cannot distinguish 5mC from 5hmC is 77.8% and is similar to previously reported in the zebrafish genome ∼70%-80% (Feng et al., 2010). However, we found that the most common configuration of 5hmC was a hemi-hydroxymethylated, hemi-methylated (5hmC/5mC) form. The ranking of methylated states is first 5mC/5mC, then 5hmC/5mC and finally the rarest form being 5hmC/5hmC. This is consistent with a random process for oxidation, leading to many hemi-hydroxymethylated sites; however the proportions do not fit a simple random model (chi-square, *p*-value < 2.2×10^-16^).

**Figure 2.**
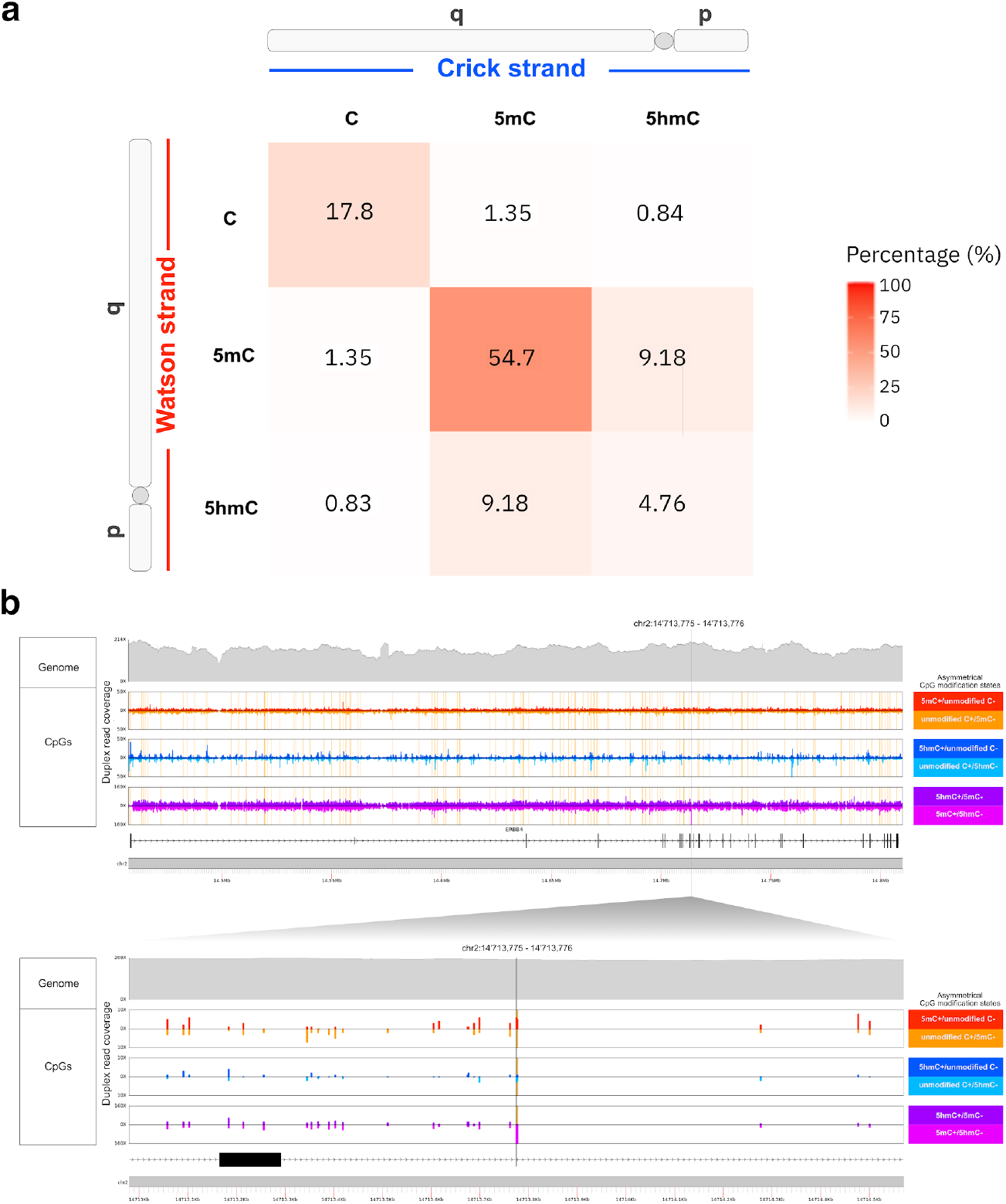
Genome-wide co-occurrence of 5hmC, 5mC and unmodified C in both strands of CpG sites in medaka’s brain tissue. (a) Heatmap depicting the frequency of modification co-occurrences in both strands of CpG sites across the entire medaka brain genome. The x-axis and y-axis depict the modification states observed in the Crick (*p* to *q* chromosome arm direction) and Watson strand (*q* to *p* chromosome arm direction) of CpG sites, respectively. (b) Genomic tracks of duplex read coverage (y-axis) in the whole *ERBB4* locus (upper panel) containing the most asymmetrically modified CpG in the genome (bottom panel, vertical grey bar). In both panels, the uppermost track (grey colour) depicts the duplex read coverage at every base-pair in the region and the other tracks (in colours) depict the duplex read coverage of each asymmetrical modification state observed at every CpG. CpGs were classified as skewed (vertical yellow bars) using the McNemar-Bowker test, when they are statistically significant for FDR adjusted *p*-values < 0.01.

As expected, when the genome is orientated with respect to the reference genome, defining the “Watson” and “Crick” strand respectively, there is no asymmetry between these strands across all CpG sites, consistent with the arbitrary orientation of the reference genome sequence. However, for each particular site we are often well-powered to see potential asymmetries at individual CpGs. We used the well established McNemar-Bowker test (a chisq statistic looking for asymmetry in a counts table) for quantifying the probability of a count asymmetry of the 6 asymmetric states. 201,863 sites, 1.4% of the CpGs in the genome were significantly skewed using a FDR correction level of 0.01; Some sites show striking asymmetries, for example with 158 out of 160 5hmC/5mC duplex reads are on the watson (p to q) direction on a CpG site inside one of the internal introns of *ERBB4* gene (Figure 2b).

### CpGs in 5’ and 3’ splice sites are preferentially hydroxymethylated in opposite strands

We explored potential reasons for the consistently skewed sites in methylation states. One candidate was the action of transcription. We hypothesized that transcribed strands could lead to skewness of modifications by imposing an intrinsic asymmetry between DNA strands due to the action of RNA polymerase or potentially co-transcriptional splicing. We recomputed asymmetry patterns orientated to the direction of transcription (Figure 3 and see methods). Overall we found that there were no large bulk asymmetries in sense vs antisense strands (Figure 3a) when averaged over all genes across their entire extent. However, when we considered individual sites we observed skewed sites that were present in both directions across all the gene features (Figure 3b). We considered skewed sites to have both passed the FDR based threshold on the McNemar-Bowker test and have an effect size of asymmetric state greater than abs(0.5) skewness - this prevents very small shifts in asymmetry at high coverage sites being noted as skewed (see methods). Although in every gene feature there are asymmetric sites in both directions, for some features there are more asymmetric sites in one direction to another - for example, around the 3’ splice sites, there are 813 compared to 408 skewed sites in the 5hmC/5mC state. To consider more subtle effects, we computed a metagene average of skewness for the 3 types of asymmetry (5mC/5C, 5hmC/5C and 5hmC/5mC) shown in Figure 3c. There is a notable increase in asymmetric 5hmC/5C in the first exon, peaking at the first intron boundary. Consistent with our per-site analysis in Figure 3b we see contrasting shifts in 5hmC/5mC between introns and exons (exons having more hydroxymethylation in the sense strand). Finally there are sharp shifts in asymmetry around the internal splice sites of genes. To explore the finer detail of splice site behaviour, we calculated the aggregate skewness per base around the 5’ and 3’ splice sites (abbreviated 5’SS and 3’SS), excluding sites which have less than 5% CpGs present. Across both splice sites the intronic side of the splice site shows more intense asymmetries, but there is not a consistent direction of asymmetry with for example two adjacent base pair positions in the 5’ SS having opposing directions of 5hmC/5mC asymmetry.

**Figure 3.**
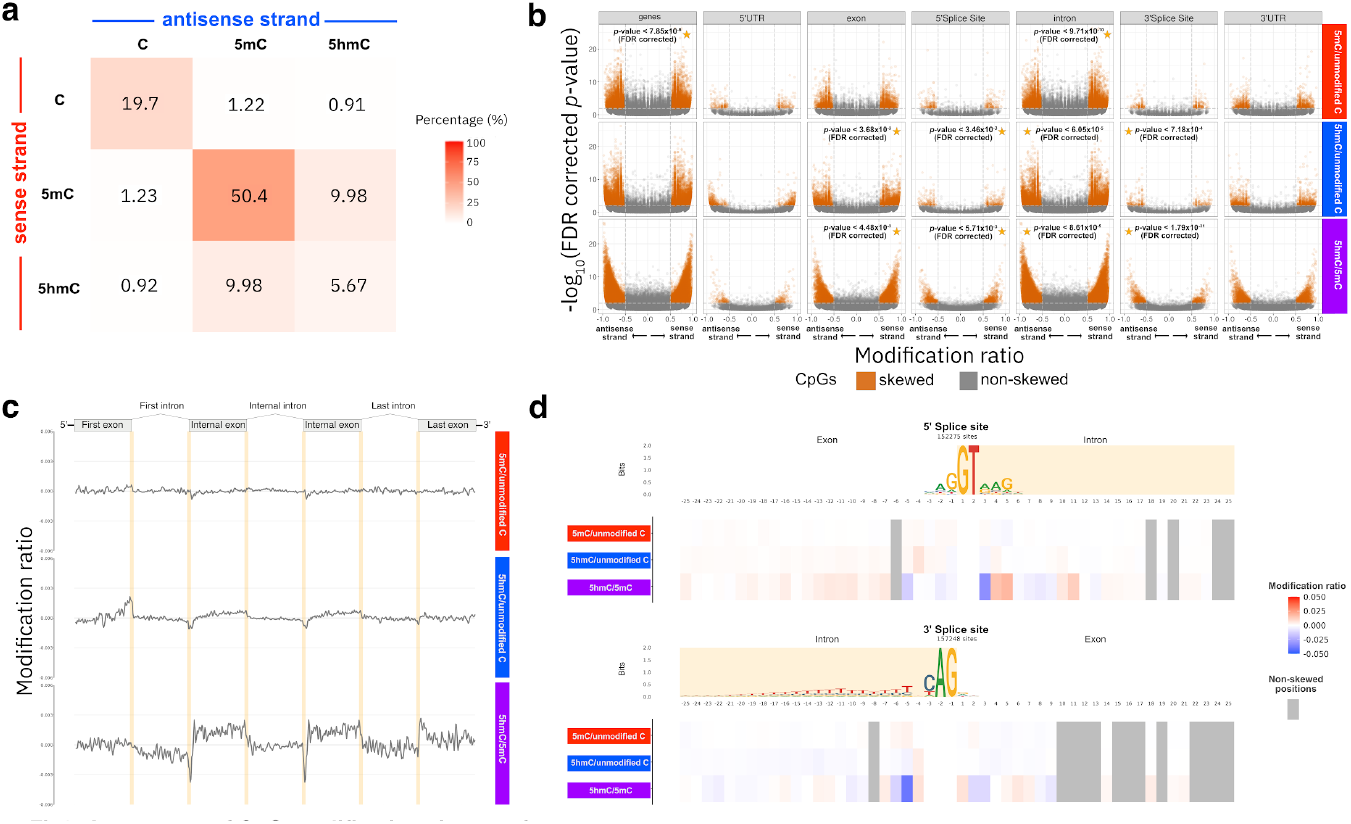
Analysis of asymmetrical states of CpGs in gene regions. (a) Heatmap depicting the frequency of modification co-occurrences in both strands of CpG sites in gene regions. The x-axis and y-axis depict the modification states observed in the antisense (blue colour) and sense strand (red colour) of CpG sites, respectively. (b) A scatterplot, inspired by the gene expression volcano plots, that depicts the ratio of difference in modification patterns between the sense and antisense (x-axis), *i*.*e*. effect size, and the associated adjusted *p*-value (y-axis) in negative logarithmic scale for each CpG. The skewed (gold colour) and non-skewed CpGs (grey colour) are depicted for the asymmetrical modification patterns (horizontal facets) across different gene features (vertical facets). CpGs were classified as skewed when they are statistically significant for FDR adjusted *p-*values < 0.01 and when showing a substantial change in effect size for modification ratios > 0.50 or < -0.50. The set of skewed CpGs with a preferential modification strand are marked with a star. (c) Metagene plot of the mean modification ratio (y-axis) of CpG asymmetric states along intron and exon features (x-axis). The intron-exon boundaries are highlighted in yellow and the internal exon of a gene is duplicated to assist with visualization of the other features. (d) 5’ and 3’ SS motifs with modification ratio values of the three asymmetrical states (heatmap), the non-significant positions are shaded in grey vertical bars.

### CpGs in families of transposable elements (TEs) are preferentially hydroxymethylated in a DNA strand

Although the gene centered analysis showed patterns of asymmetries relative to transcription, this was not an adequate explanation for the large number of asymmetric sites across the genome. We considered features such as telomere or centromere association, low complexity regions, overall CpG content and transposable elements (TEs). For the latter class, we saw that some dispersed transposable element families have a large number of skewed sites often in a consistent configuration relative to the transposon orientation. We observed that 26 families of TEs have CpGs with 5hmC preferentially occurring in one strand of the DNA duplex (Figure 4a) and with strand preference being independent for each TE family (Figure 4b). Overall 17% of skew sites can be explained by their presence in a TE family, with an additional 15% present in exons or in a splice site. Of the remaining sites, 36% are in introns but not near splice sites or in a TE and 31% in intergenic regions but not in a TE.

**Figure 4.**
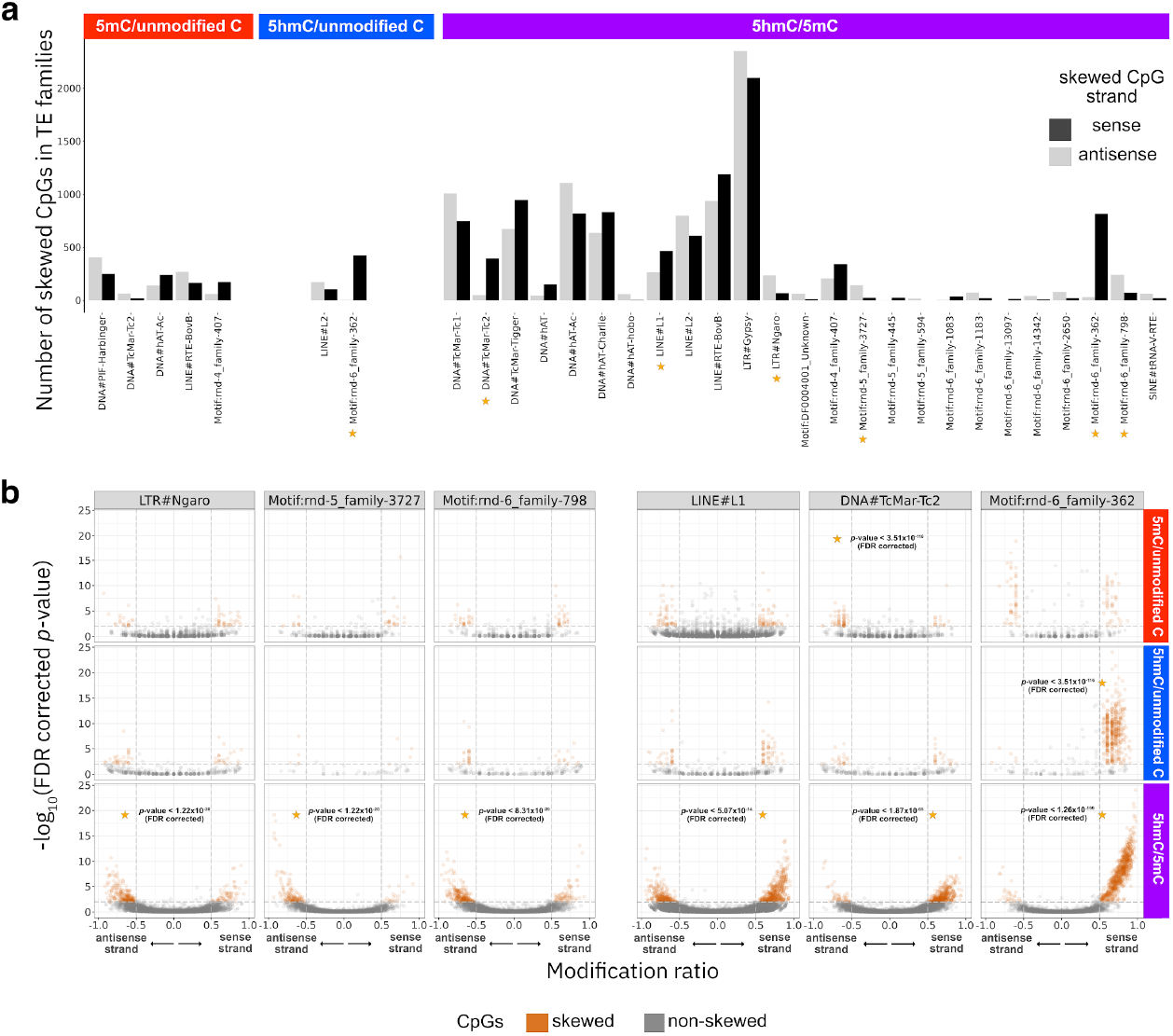
Analysis of asymmetrical states of CpGs in skewed TE families. (a) Frequency of skewed CpGs (y-axis) in the sense (black colour) and antisense strand (grey colour) of 26 TE families (x-axis) with preferential modification patterns in a DNA strand. The top skewed TE families are marked with a star along the different asymmetrical modification patterns (vertical facets). (b) A scatterplot, similar to Figure 3b depicts the ratio of difference in modification patterns between the sense and antisense (x-axis), *i*.*e*. effect size, and the associated adjusted *p*-value (y-axis) in negative logarithmic scale for each CpG. The skewed (gold colour) and non-skewed CpGs (grey colour) are depicted for the asymmetrical modification patterns (horizontal facets) across different TE families (vertical facets). CpGs were classified as skewed when they are statistically significant for FDR adjusted *p-*values < 0.01 and when showing a substantial change in effect size for modification ratios > 0.50 or < -0.50. The set of skewed CpGs with a preferential modification strand are marked with a star.

## DISCUSSION

In this work we have explored the landscape of CpG modifications both considering methylation and hydroxymethylation and considering the co-occurrence of modifications on the same double stranded DNA molecule. Although there has been extensive previous work on exploring hydroxymethylation (He et al.,2021; Cui et al., 2020) and individual sites have shown asymmetric hemi-methylation patterns (Xiong et al., 2024; Hua et al., 2024; C. Xu & Corces, 2018) we believe we have performed the most comprehensive analysis of CpG modifications in a strand aware manner. This is due to the Oxford Nanopore technology being able to natively distinguish methylation and hydroxymethylation and that a proportion of Oxford nanopore reads are sequenced in a duplex format. It is worth highlighting that the most common way to assess methylation, which is bisulphite treatment followed by sequencing or array read out, assesses the union of methylation and hydroxymethylation.

Our overall statistics are reasonably consistent with previous work; Wen et al., 2014 reported 13.4% of CpG’s being highly hydroxymethylated in mouse neuronal tissues. Our duplex based analysis does not have a 1 to 1 comparison with the simplex based bisulphite and TET enzyme readout. We observed 25% of duplex bases crossing CpGs as having hydroxymethylation. A surprise is that the majority of hydroxymethylation is found in a hemi-hydroxymethyl/hemi-methylation format. This finding is highly congruent with previously reported results from single-molecule fluorescence assays (Song et al., 2016), which only looked at subset of sites and had a wide confidence interval. This hemi hydroxymethylation is somewhat consistent with a largely random process, but also provides the opportunity to assess whether there is a strand bias of hydroxymethylation. For 1.4% of CpG sites we see a statistically significant skewness of the 5hmC/5mC modifications. Having explored different possibilities for this skewness, two genomic features are associated with skewed behaviour. The most impactful are TEs which account for 17% of the skewed sites. There is no specific direction of the skewness relative to TE transcription. The other association is with transcribed genes, with a bias in skewed direction between introns and exons, and a notable pattern of skewed sites near splice sites. As methylation and hydroxymethylation are properties of DNA, but splicing is an RNA phenomena this implies that there is co-transcriptional splicing which impacts the accessibility or activity of DNA methylation processing, in particular its oxidation to hydroxymethylation. An alternative hypothesis is that hemi-hydroxymethylation is an actively maintained mark to help designate splice sites but this seems far less likely from a mechanistic perspective, noting that many pre-mRNAs can be accurately spliced in the absence of DNA. Either way, hemi hydroxymethylation is a potential new biomarker for splicing behaviour.

The presence of a substantial number of skewed hemi-hydroxymethylation in TEs also requires explanation. Our proposed model is that many TEs have evolved to actively demethylate, removing this largely repressive mark in an arms race with the host somatic genome, which is actively methylating. In this arms race it would be natural for TEs to try to co-opt the TET enzymes for oxidation of CpG methylation, leading to eventual derepression. As there is a known large change in the regulatory landscape in neuronal tissues, including methylation patterns and active demethylation patterns, it follows that we see different patterns of active methylation and demethylation between the host and TE genomes, giving rise to a patchwork of different active demethylation patterns, including hydroxymethylation.

Whatever the cause of these signals - whether co-transcriptional splicing, transcriptional or TE based - hydroxymethylation and hemi-hydroxymethylation is a potential new genome-wide biomarker of regulatory status. This has potential in neuronal cancers where CpG methylation is already useful for cancer classification (Benfatto et al., 2025; Brändl et al., 2025; Vermeulen et al., 2023). It will be interesting to explore the hemi-hydroxymethylation signal in the context of neuronal tissue cancers.

More generally, the ability to accurately call DNA modifications in a strand specific manner provides for a richer understanding of the native DNA formulation, using technologies such as ONT. Other technologies, such as PacBio, provide read out for certain modifications. These modifications in their double stranded context provides a readout of both the true native state of DNA in these cells, which potentially impacts transcription factor binding and other processes, and also provides a novel way to read out the state of the cell beyond “just” methylation. Our current study has utilised the smaller genome of the medaka fish and our serendipitous choice of neuronal tissue as a source of DNA to allow for a comprehensive genome wide view; however, it is entirely straightforward to apply to mammalian and human samples where native DNA is available.

## ACKNOWLEDGEMENTS

This project has received funding from the European Research Council (ERC) under the European Union’s Horizon 2020 research and innovation programme (grant agreement No 810172). W.S.G., T.F. and E.B. are funded by the European Molecular Biology Laboratory (EMBL). We are thankful to Charlotte West and Nicola de Maio from the Goldman lab at EMBL-EBI for useful discussions on the analysis of TEs. We also thank Amos Pierroti, for kindly providing the medaka illustrations that were used in the manuscript.

## CONTRIBUTIONS

Conception of the project: E.B; Project management and supervision: E.B., J.W., T.F.; Inbreeding of fishes: F.L.; Sample preparation: F.L.; Data analysis: W.S.G.; Manuscript writing: E.B., T.F., W.S.G. All author(s) read and approved the final manuscript.

## MATERIALS AND METHODS

### Nanopore sequencing of *Oryzias latipes* genomes

Brain tissues were dissected from 50 *Oryzias latipes* (medaka) individuals from the Medaka Inbred Kiyosu-Karlsruhe (MIKK) Panel (Leger et al., 2022). In brief, DNA was extracted from brain tissues and prepared for ligation with sequencing Kit V14. Eight samples were multiplexed using native barcoding Kit 24 V14 (SQK-NBD114-24) and the remaining 42 were multiplexed using native barcoding Kit 96 V14 (SQK-NBD114-96). With the exception of eight samples that were multiplexed in groups of four, all samples were multiplexed in groups of three and loaded for sequencing in R10.4.1 flow-cells (FLO-PRO114M) on a PromethION 24 instrument. Sample sequencing was performed selecting an average speed of 400bps per second and at a sampling rate of 4kHz or 5kHz.

### Base and modification calling of Nanopore sequencing reads

#### Simplex read calling

Demultiplexed raw FAST5 files generated by the nanopore sequencing device were converted to POD5 file format using ONT *pod5 convert fast5* v0.1.5. The two sets of “Pass” and “Fail” POD5 files for each sample were merged in a single POD5 file using *pod5 merge* v0.1.5. Then, POD5 files were used for simplex base and modification calling of 5mC and 5hmC using the super-accurate basecalling model “dna_r10.4.1_e8.2_400bps_sup@v4.1.0“ with ONT *dorado basecaller* v0.3.0, and parameters “--emit-moves --device ‘cuda:all’ --modified-bases ‘5mCG_5hmCG’“.

#### Duplex read calling

Simplex reads called in modBAM format were used to detect pairs of complementary simplex reads belonging to the same double-stranded DNA molecule using ONT *duplex_tools pair* v0.3.1 with default parameters. To basecall duplex reads in BAM format, the list of read ID pairs and the source POD5 file of the simplex reads for each sample was provided to ONT *dorado duplex* v0.3.0 using parameters “--emit-sam” and the super-accurate basecalling model “dna_r10.4.1_e8.2_400bps_sup@v4.1.0“.

#### Duplex read calling from chimeric reads

To identify simplex chimeric reads and split each raw read signal in two, ONT *duplex_tools split_pairs* v0.3.0 was used with parameter ‘--threads 5’. For each sample, the simplex reads in modBAM format and the source POD5 file of each simplex read set were provided as input. The resulting split read signal in POD5 format and the list of read ID pairs were passed to *dorado duplex* for duplex basecalling using the aforementioned parameters. BAM files of duplex reads without modifications were obtained as output, at the moment *dorado* is unable to call modifications in duplex reads.

#### Duplex modification calling

5mC and 5hmC modifications were called in the main set of duplex reads and the duplexes called from chimeric reads, using ONT *remora infer duplex_from_pod5_and_bam* v2.1.2 with default parameters. The source POD5 file, the modBAM file and the list of read ID pairs of the simplex reads, as well as the duplex read BAM file were used as input to *remora* and providing the super-accurate modification model “dna_r10.4.1_e8.2_4khz_400bps_sup_v4.1.0_5hmc_5mc_CG_v2.pt”.

### Processing of duplex read sequences and alignment

#### Filtering of duplex read sequences

After calling modifications in the two sets of duplex reads for each sample, we discarded duplex reads in the two sets that had the same simplex read ID associated to more than one duplex read. In extremely rare occasions a simplex read can be identified as chimeric, and in parallel paired to another simplex read in the main duplex read set. The paired simplex reads from the discarded duplex reads were treated as simplex for downstream analysis. After filtering the two set of duplex reads, the modBAMs of each reads set were pooled using *samtools merge* v1.17. Finally, reads smaller than 50 bases or with mean quality of less than 10 were discarded with pysam v0.19.0 (https://github.com/pysam-developers/pysam).

#### Alignment and alignment filtering

Reads in modBAMs were transiently converted to FASTQ format and preserving all read tags with *samtools fastq* v1.17, using the parameter “-T ‘*’“. The FASTQ reads were piped to Minimap2 v2.24-r1122 (Li, 2018) for alignment to the medaka reference genome (*HdrR* ASM223467v1) from the ensemble release 99 (Cunningham et al., 2022) using the parameters “-x map-ont -a -t 30 -y --secondary=no –MD”. Reads with secondary or supplementary alignments were discarded, as well as unmapped reads, using *samtools view* v1.17. modBAM files were sorted by coordinate and indexed with *samtools sort* and *samtools index* v1.17 for downstream analyses.

### Identification of CpG sites with skewed modification patterns between Watson and Crick strands

*Extraction of modifications on CpG nucleotides from duplex reads*. To obtain the modification patterns on both strands of CpG nucleotides in duplex reads, we first calculated the coordinates in BED format of all CpG nucleotides in the medaka genome using ONT *modkit motif-bed* v0.1.11 with parameters “CG 0”. Then, we used ONT *modkit extract* v0.1.11 with default parameters to extract the genomic coordinates and probabilities of 5mC and 5hmC modifications in both strands of the duplex reads, but restricting the output to the coordinates of the CpG nucleotides from the previously produced BED file.

#### Assignment of CpG states

Raw modification probabilities were converted from their original discrete scale [0,255] to a continuous scale [0,1]. As probabilities of observing canonical nucleotides (*e*.*g*. cytosines) are not explicitly encoded in modBAMs, but only modified states, we computed the probability of observing the unmodified state of cytosine as

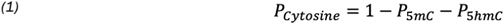

The modification state (*i.e*. unmodified C, 5mC or 5hmC) of a cytosine in a single strand was assigned to the state with the largest probability value. Then, the modification states of the two cytosines occurring in both strands of the palindromic CpG nucleotides, were used to assign one of nine possible states (*i.e*. 5mC+/unmodified C-, 5mC+/5mC-,5mC+/5hmC-, 5hmC+/unmodified C-, etc.) to each CpG found in a single duplex read. Then, for each medaka sample all the occurrences of CpG states across all reads were counted to create a matrix of dimensions *N* x 9, where *N* is equal to the number of CpGs in the medaka genome.

### Identification of CpG sites with skewed modification patterns between DNA strands

#### Statistical analysis of modifications in individual CpG sites

All the count matrices of the samples were summed to create a single count matrix. CpG sites with low or high coverage (*i.e*. the first 15 percentiles and the last 10 percentiles of the coverage distribution) were removed from downstream analyses. To identify the CpG sites with modification skewness between Watson and Crick strands, we used the McNemar-Bowker to test for symmetry of counts in the modification patterns observed between the two DNA strands. The McNemar-Bowker test uses a chi-square distribution. In our case, we computed the chi-square value with three degrees of freedom as defined below:

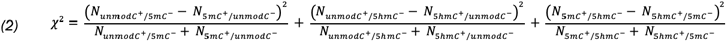

Where *N* is the number of occurrences of one of six asymmetrical modification states, with strand orientation, observed at each CpG.

As zero counts for some modification patterns can occur in some CpGs, we added a pseudo count of one to all counts before computing the McNemar-Bowker test. After performing the statistical test, *p*-values were corrected for multiple testing using the false discovery rate (FDR) method.

The modification ratio for each of the three asymmetrical modification states at a single CpG was defined as:

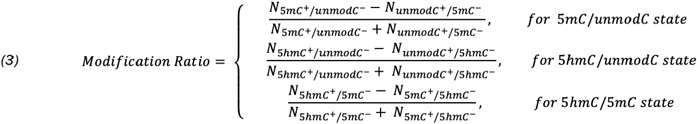

Finally, CpG sites were reported as skewed when they were statistically significant for adjusted *p*-value < 1×10^-2^ and when showing a substantial change in effect size for modification ratio > 0.50 or modification ratio < -0.50.

### Identification of CpG sites with skewed modification patterns between DNA strands in gene features

#### Processing of medaka gene feature annotations

The medaka gene annotations from the ensemble release 109 (Harrison et al., 2024) were retrieved and processed in R v4.4.1 (R Core Team, 2024). In brief, medaka annotations containing transcript, exon, 5’ and 3’UTR features were read using *makeTxDbFromGFF*, and intron intervals for each transcript were calculated using the *intronsByTranscript* function from the *GenomicFeatures* library v1.57 (Lawrence et al., 2013) . The 5’ and 3’ Splice Site features were defined as the -/+10 bp interval from the 5’ or 3’ exon-intron boundary. Only transcript features from the Ensembl Canonical transcript collection were retained for downstream analysis.

#### *Statistical analysis of modifications on individual CpG sites* in gene features

Genomic coordinates of CpG sites were intersected with the previously processed medaka gene features using *bedtools intersect v2.30.0* (Quinlan & Hall, 2010). CpG sites overlapping gene annotations were retained and nucleotide modification strands of each CpG were re-oriented to match the sense and anti-sense strand of the gene where the CpG is located. To mitigate annotation redundancy when a CpG overlapped multiple gene features, a random annotation was selected. Then, the McNemar-Bowker test was used (see *Statistical analysis of modifications in individual CpG sites* section) to test the modification skewness between sense and anti-sense strands at each CpG site.

### Metagene analysis of asymmetrical modification states in the medaka genome

#### Processing of medaka metagene feature annotations

The previously processed medaka gene annotations (see *Processing of medaka gene feature annotations*) were filtered to retain only intron and exon features of transcript models with at least one intron. A metagene label was assigned to intron and exons according to their position in the transcript model. Namely, (i) the first exon and intron were labeled as “First exon” and “First intron”, respectively; (ii) the last exon and intron were labeled as “Last exon” and “Last intron”, respectively, and (iii) all the remaining internal exons and introns were labeled as “Internal exon” and “Internal intron”, when present in a transcript model.

After grouping the intron and exon in metagene features (*e.g*. First exon, First intron, etc.), the smallest features were discarded before binning them. In brief, the length of gene features was computed (*ℓ*), and the 1st percentile (*i.e*. 75 bps) of the length distribution was used as threshold to filter out all gene features with ℓ *<* 75 bps. The remaining gene features were segmented in 75 contiguous bins, and the genomics intervals of the features were used for intersection with CpG coordinates.

#### Computation of modification ratios of asymmetrical modification states

Firstly, we added a pseudo count of one to all nine modification pattern counts observed at every CpG in the 50 independent count matrices (*i.e*. one matrix per medaka individual) of dimensions *N* x 9 (see *Assignment of CpG states* section). Then, the modification ratio for each of the three asymmetrical modifications (*i.e*. 5mC/unmodified C, 5hmC/unmodified C and 5hmC/5mC) was computed for all CpGs in the 50 medaka individuals as defined in equation (3). When a CpG was not observed in a medaka individual, a modification ratio of zero was registered for the three asymmetrical modification states. Then, three mean modification ratios (*i.e*. one per asymmetrical modification state) were computed for each CpG across the 50 samples, to create a single matrix of dimensions *N* x 3, where rows are equal to the number of CpGs in the medaka genome and the columns to the number of mean modification ratios.

The genomic coordinates of each CpG with their mean modification ratios were intersected with the previously binned medaka metagene features (see *Processing of medaka metagene feature annotations* section) using *bedtools intersect v2.30.0*. CpG sites overlapping gene annotations were retained and the signs of the mean modification ratios for each CpG were re-oriented to match the sense and anti-sense strand of the gene feature where the CpG is located.

Two sets of mean modification ratios were computed. Firstly, the set of mean modification ratios of a single bin was computed and defined as the three mean modification ratios (*i.e*. one per asymmetrical modification state) of all CpGs contained in the *i*-th bin of a gene feature. Then, the set of mean modification ratios of a bin was used to compute the set of mean modification ratios of the *i*-th bin in a metagene feature group (*e.g*. First Exon, First Intron, Internal Exon, etc.).

### Analysis of asymmetrical CpG modification states in 5’ and 3’ SSs motifs

#### Creation of 5’ and 3’ SSs motifs

To create the 5’ and 3’ SSs motifs, 50-bp coordinates around the 5’ and 3’ exon-intron boundaries were created in R, and they were used to extract the corresponding fasta sequences in the medaka reference genome using *bedtools getfasta* v2.30.0 with parameters “-s -name”. The retrieved fasta sequences were then processed in R with Biostrings v2.74.0, and only the sequences containing canonical splice sites, *i.e*. GT and AG at the 5’ and 3’ SSs respectively were retained. Finally, the Position-Specific Scoring Matrices (PSSMs) were computed using Biostrings for the 5’ and 3’ SSs set of fasta sequences to create two independent motifs.

#### Selection of CpGs in 5’ and 3’ SS motifs

CpG coordinates were intersected with the 50-bp coordinates around the 5’ and 3’ exon-intron boundaries (see *Creation of 5’ and 3’ SSs motifs* section) with bedtools intersect v2.30.0 and parameters “-wao”. Only CpGs overlapping exon-intron boundaries were retained and an index was assigned to each CpG according to their position in the 50-bp exon-intron boundary.

#### Computation of modification ratios of asymmetrical modification states in 5’ and 3’ SSs

The nucleotide modification strands of each CpG were re-oriented to match the sense and anti-sense strand of the exon-intron boundary where the 5’ and 3’ SSs were contained. Then, the set of mean modification ratios (*i.e*. one per asymmetrical modification state, see *Computation of modification ratios of asymmetrical modification states*) was computed for all CpGs at each index position in the 5’ and 3’ SS motifs.

#### Identification of positions at 5’ and 3’ SSs motifs with skewed modification states

To identify skewed modification states at each index position in the 5’ and 3’ SS motifs, the modification pattern counts of all CpGs were summed at each position in the 5’ and 3’ SS motifs. Then, 100 contingency matrices (*i.e*. 50 for 5’ and 50 for 3’ SSs) of 3x3 dimensions were created to summarise the nine possible modification pattern counts of CpGs at each position in the 5’ and 3’ SS motifs. A pseudo count of one was added to every modification pattern count in the 100 contingency matrices and the McNemar-Bowker test was performed on each matrix. The resulting *p*-values of each test were then corrected for multiple testing using the FDR method. Finally, positions in the 5’ and 3’ SSs with a corresponding *p*-value <0.01 were identified as significantly skewed.

### Identification of CpG sites with skewed modification patterns in transposable elements

#### Processing of medaka annotations of Tes

The annotations of repetitive elements in the medaka genome were retrieved from Fitzgerald et al., 2022. Simple repeats and low complex regions were discarded, and only TE annotations were retained for downstream analyses.

#### Statistical analysis of modifications on individual CpG sites in Tes

Genomic coordinates of CpG sites were intersected with the previously filtered TE annotations using *bedtools intersect v2.30.0*. CpG sites overlapping TE annotations were retained and nucleotide modification strands of each CpG were re-oriented to match the sense and anti-sense strand of the TEs where the CpG is located. To mitigate annotation redundancy when a CpG overlapped multiple TE annotations, a random annotation was selected. Then, the McNemar-Bowker test was used, as described above (see *Statistical analysis of modifications in individual CpG sites* section), to test the modification skewness between sense and anti-sense strands at each CpG site.

## SUPPLEMENTARY FIGURES

**Supplementary Figure 1.**
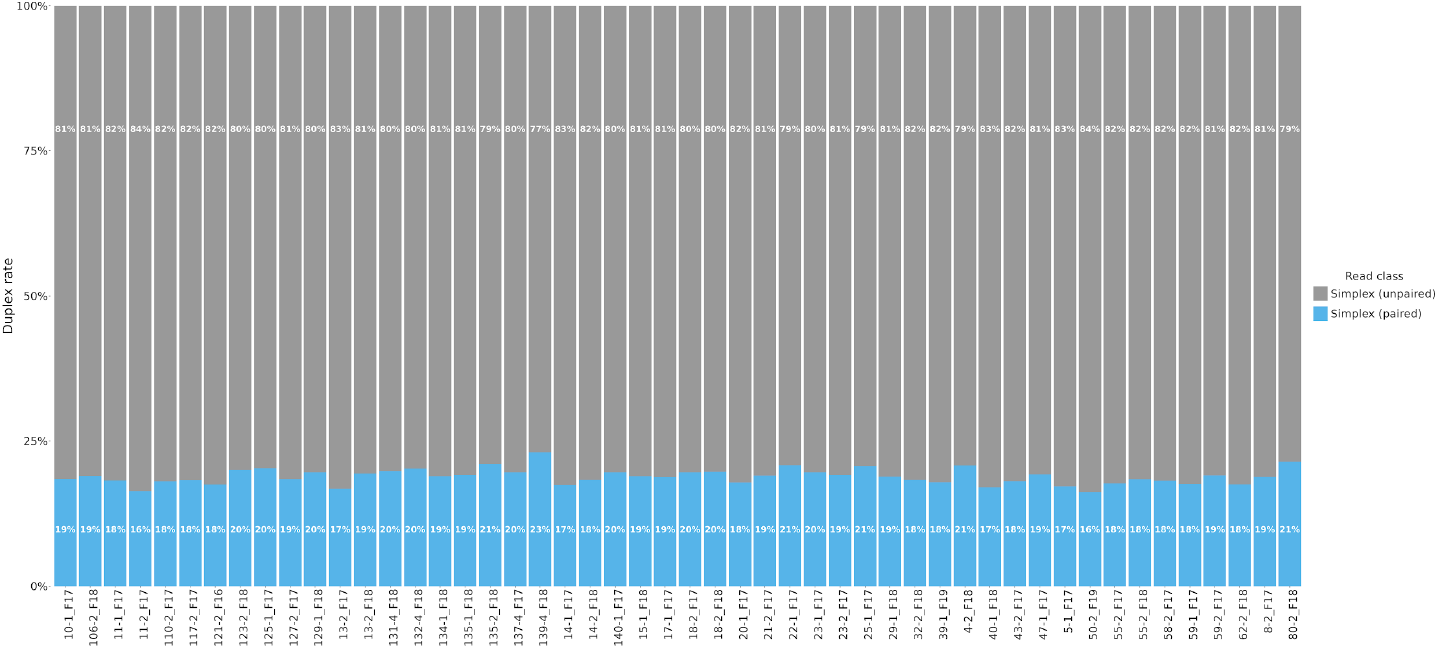
Duplex read yield in medaka samples. The percentage of simplex reads (y-axis) paired in a duplex read form (blue colour) and the unpaired simplex reads (gray colour) are depicted for the 50 medaka samples (x-axis).

